# Outer Membrane Vesicles (OMVs) Produced by Plant Beneficial Rhizospheric Bacteria Enter Root Epidermal Cells

**DOI:** 10.1101/2025.07.18.665571

**Authors:** Sofija Nešić, Dragana Bosnić, Aleksandra Divac Rankov, Maja Kosanović, Jelena Samardžić, Bojana Banović Đeri, Dragana Nikolić

## Abstract

Extracellular vesicles (EVs) are shown to play an important role in the interaction between bacteria and human cells, as well as between fungi and plants; however, much less is known about their role in plant-bacteria relations. Outer membrane vesicles (OMVs), EVs produced by Gram-negative bacteria, were shown to have both immuno-activating and immuno-suppressive effects on plants, and a direct interaction of OMVs with plant cell membranes was shown for one phytopathogenic bacterial strain. Data on OMVs’ participation in the symbiotic relationship between plant-beneficial bacteria and plants are even more limited, and are mainly related to the rhizobia-legume interaction.

To investigate the interaction of OMVs of non-rhizobial phytobeneficial bacteria with plant cells, we isolated and characterized OMVs produced by *Paraburkholderia phytofirmans* PsJN, a strain well known for its growth-promoting and stress-protective effects in a wide range of plant species. Using OMVs labelled with fluorescent lipid-binding dye Vybrant^TM^ DiD and confocal laser scanning microscopy, we showed that PsJN-OMVs interact directly with the root hairs and epidermal cells of *Arabidopsis thaliana* and tomato. Moreover, by employing octadecyl-rhodamine B chloride (R18), we demonstrated that PsJN-OMVs can fuse with plant cell membranes.

The findings presented in our study are an important step in elucidating the potential role of OMVs in the early stages of root colonization and the establishment of symbiotic relationships. Increasing our understanding in this area could pave the way for the use of these nano-vehicles to develop eco-friendly solutions for overcoming environmental challenges and supporting sustainable agriculture.

## Introduction

Plant growth-promoting bacteria (PGPB) enhance plant growth, productivity, and stress resilience through nitrogen fixation, siderophore production, phosphate solubilization, and phytohormone production or reduction. They also help plants combat pathogens through induced systemic resistance (ISR), as well as biocontrol activity by producing antibiotics and antifungal compounds (Backer et al., 2018; Olanrewaju et al., 2017). In exchange, plants supply the bacteria with reduced carbon compounds and other metabolites. Root exudates serve as key recruitment signals for bacteria, while bacterial features, such as chemotaxis and biofilm formation, facilitate effective root colonization. To colonize roots, PGPB must evade the plant immune system. In other words, plant defense responses are finely tuned to distinguish mutualists from pathogens (Zhang & Kong, 2022). Also, in the rhizosphere, bacteria compete with other microorganisms for nutrients and niches.

These complex networks of inter-species interactions require multifaceted exchange of signal and effector molecules. Among other ways of signal conveyance, extracellular vesicles (EVs) are recognized as an important tool for intercellular, inter-organismal, and even inter-kingdom communication (Cai et al., 2018; Weiberg et al., 2013; Zhao et al., 2025). These nanoscale structures, which are secreted by all eukaryotic and prokaryotic cells, carry cellular components such as lipids, proteins, nucleic acids, and metabolites. What distinguishes them and gives them an advantage over other secretion systems is that EVs deliver signals at a distance, without direct contact between a donor and a recipient cell. Moreover, EVs envelop their cargo in a lipid membrane, thereby protecting it and allowing targeted delivery of high concentrations of active substances to the recipient cells (Bomberger et al., 2009). Both EV membrane and cargo molecules are shown not to be merely randomly released cellular components, but rather a result of selective packaging. EVs’ protein, lipid, and RNA compositions overlap with the intracellular content, but some elements are found to be enriched in EVs or predominantly retained inside the cells (Malabirade et al., 2018; Orench-Rivera & Kuehn, 2021). The molecular mechanisms underpinning selective loading of eukaryotic EVs are partially described, while those responsible for bacterial vesicles remain poorly understood. Analyses of the enriched EVs’ content indicated that EV secretion serves as a means for eliminating damaged components, allowing for a fast response and stress resilience (McBroom & Kuehn, 2007; Schwechheimer & Kuehn, 2013). However, the packaging of specific compounds such as toxins in pathogenic EVs has also been recognized (Kesty et al., 2004; Rivera et al., 2010). Regarding nucleic acids, it is suggested that EVs may serve both to remove unnecessary cellular RNAs and to transfer RNAs that could have signaling functions (Dellar et al., 2024).

EVs may contribute to many vital processes in bacteria. In communication with other bacteria within the population, bacteria transfer signaling substances like quorum sensing molecules in EVs to help coordinate activities of population members, for instance, during biofilm formation (Mashburn & Whiteley, 2005; Schooling & Beveridge, 2006). They are also shown to facilitate horizontal gene transfer and aid in nutrition (C. Li et al., 2022; Lin et al., 2017). To outcompete other microorganisms, bacteria use EVs as one of their weapons, sending them filled with antibiotics to their competitors or enemies (Ñahui Palomino et al., 2021).

The most intriguing aspect of bacterial EVs’ action is in communication with eukaryotic host cells. Both pathogenic and commensal bacteria are shown to send signals to their hosts, carried by EVs. Pathogenic EVs are known to transfer virulence factors, which can include proteins as well as small RNAs that target host defense genes (Choi et al., 2017; Kesty et al., 2004). The research on the role of EVs in the interaction between bacterial and human cells and the mechanisms of EVs’ entry into mammalian cells has received much attention lately, especially regarding outer membrane vesicles (OMVs) produced by Gram-negative bacteria. Moreover, extensive efforts have been made to harness OMVs for the development of vaccines and drug delivery systems (Jiménez‐Guerrero et al., 2023; Qing et al., 2019). However, little is known about OMVs secreted by plant-associated bacteria and their role in plant-bacteria interactions. Both immuno-activating and immuno-suppressive effects of OMVs produced by different pathogenic bacteria on plants have been demonstrated (Bahar et al., 2016; Janda et al., 2023; McMillan et al., 2021; Wu et al., 2024). Nonetheless, a direct interaction of OMVs with plant cell membrane was shown only for one phytopathogenic bacterial strain (Tran et al., 2022). Information on the involvement of OMVs in the symbiotic relationship between plant-beneficial bacteria and plants is even more limited and mainly related to rhizobia-legume interaction (Ayala-García et al., 2025; D. Li et al., 2022; Taboada et al., 2019).

In this study, we have characterized OMVs secreted by a non-rhizobial plant-beneficial rhizospheric bacterium, *Paraburkholderia phytofirmans* PsJN, well known for its plant growth-promoting properties and stress protection in many plant species (Esmaeel et al., 2018). Further, we explored whether PsJN-OMVs interact with root epidermal cells. The interaction was tested in two plant species: the widely utilized model plant *Arabidopsis thaliana* and the worldwide-grown crop tomato. Previously, it has been demonstrated that PsJN promotes the growth of both *A. thaliana* and tomato, and alleviates abiotic stresses such as heat and salt stress (Issa et al., 2018; Ledger et al., 2016; Macabuhay et al., 2022). Improved tolerance to biotic stresses and priming of plant immune responses were also shown in *A. thaliana* (Nešić et al., 2025; Timmermann et al., 2017). Although PGPBs are recognized as a promising sustainable alternative to conventional agriculture, a comprehensive understanding of the molecular mechanisms underlying beneficial bacteria-plant interactions is still missing. The successful colonization and establishment of the symbiotic bacteria-plant relationship require a multifaceted exchange of signals. Considering the role of EVs in other inter-kingdom microorganism-host interactions, we address the question about the potential participation of OMVs in the early stages of root colonization by non-rhizobial PGPB.

## Materials and Methods

### Bacterial strain and growth conditions

*Paraburkholderia phytofirmans* PsJN (obtained from Leibniz Institute DSMZ - German Collection of Microorganisms and Cell Cultures) was cultured at 30°C in an orbital shaker, in mineral medium MSM, containing in [mM]: 13.03 K_2_HPO_4_, 6.98 KH_2_PO_4_, 5.07 (NH_4_)_2_SO_4_; and in [µM]: 34.2 Na_2_EDTA, 6.96 ZnSO_4_, 6.8 CaCl_2_, 18 FeSO_4_, 0.826 NaMoO_4_, 0.8 CuSO_4_, 1.682 CoCl_2_, 6.16 MnSO_4_ and 487 MgSO_4_. Glucose (0.2%) was added as a carbon source.

### Plant Material and Growth Conditions

Seeds of *Arabidopsis thaliana* Col-0 or tomato (*Solanum lycopersicum*, Novosadski jabucar) were surface-sterilized with 70% (v/v) ethanol for 5 min, followed by 20 min incubation in a sterilizing solution (10% commercial bleach, 0.05% (v/v) Tween 20, dH_2_O). The seeds were washed 3 times with sterile distilled water (dH_2_O). After 5 days of stratification in the dark at +4 °C, the seeds were sown under sterile conditions on half-strength MS medium (Murashige & Skoog, 1962) supplemented with sucrose and grown vertically. Seedlings of *A. thaliana* were grown at 21 °C under short-day conditions (10 h day /14h night) at a light intensity of 150 μmol m^−2^s^−1^ and a relative humidity of 55-65 %. Tomato seedlings were germinated at 24 °C for 3 days in the dark and then transferred to a short-day light regime (10 h day/14 h night) with a light intensity of 200 μmol m^−2^s^−1^ and a relative humidity of 55-65%. Four-day-old seedlings were used for OMV treatment.

### Isolation and purification of OMVs secreted by *P. phytofirmans* PsJN

PsJN was cultured in MSM medium until an optical density (OD) of 1 (mid-logarithmic phase) was reached. The bacterial cells were pelleted at +4°C/6000g/20 min. The supernatant was centrifuged again under the same conditions and gently filtered through microporous membranes (0.45 and 0.22 µm) to remove all remaining bacterial cells and cellular debris. OMVs were isolated from the filtered supernatant using an affinity-based column system, highly selective for specific enrichment of OMVs, the ExoBacteria™ OMV Isolation Kit (System Biosciences, EXOBAC100A-1), according to the manufacturer’s instructions. The buffer of the isolates was exchanged by dialysis (dialysis tubing cellulose membrane molecular weight cut-off 14,000) in PBS (phosphate buffer saline).

### Determination of concentration and size of particles by Nanoparticle Tracking Analysis (NTA)

The concentration and size distribution of the particles were analyzed by nanoparticle tracking analysis (NTA) using the ZetaView Quatt PMX-430 instrument and ZetaView software version 8.05.16 SP3 (Particle Metrix, Inning, Germany). Calibration of the instrument, including automatic cell verification, camera and laser alignment, and focus optimization, was performed according to the manufacturer’s protocol using 100 nm polystyrene beads. Prior to analysis, samples were diluted 1:250 in PBS to ensure that the particle concentration was within the optimal detection range. Particle illumination was achieved using a blue 488 nm laser. For measurements in scattering mode, video acquisition was set to a sensitivity of 78, a shutter speed of 100, and a frame rate of 30 frames per second, with recordings made over a single measurement cycle. A minimum area of 10, a maximum area of 1000, and a minimum brightness threshold of 30 were set for data processing. Each sample was analyzed at 11 distinct positions to calculate the median particle size.

NTA analysis of octadecyl-rhodamine B chloride (R18) -labeled OMVs was performed using a 520 nm laser and a 550 nm filter, a sample dilution of 1:67, and an instrument setting of sensitivity 90 and shutter speed 200.

### Quantification of lipopolysaccharide (LPS), proteins, and RNA

LPS concentration in the samples and control was measured using the Thermo Scientific Pierce™ Chromogenic Endotoxin Quant Kit according to the manufacturer’s instructions. Protein concentration in OMV isolates was determined by Pierce™ BCA Protein Assay Kit. RNA from OMVs was isolated using the EVery EV RNA Isolation Kit (System Biosciences), and RNA concentration was measured using the Qubit™ RNA HS (High Sensitivity) Assay Kit, following the manufacturer’s protocol.

### Labeling of OMVs and removal of unbound dye

Following buffer exchange to PBS, the isolated OMVs (2.4 mL) were labeled with the lipid-binding fluorescent dye Vybrant™ DiD Cell-Labeling Solution (Invitrogen™) at 10 µM/37°C/1h. To remove unbound DiD dye, DiD-labeled OMVs were purified by ultracentrifugation in a discontinuous iodixanol (OptiPrep™) density gradient. Two different gradients were tested: one with layers ranging from 21% to 35% and the other, with finer layer density differences, from 21% to 28%. The finer density gradient ensured consistency and reproducibility in completely removing the unbound DiD dye, as indicated by the control (data not shown); thus, it was used for washing DiD-OMVs and the control prior to application to the roots. DiD-OMVs or the control were mixed with OptiPrep™ to obtain 28% iodixanol, which was loaded at the bottom of the gradient, below the layers: 21%, 22%, 23%, 24%, 25%, and 26%. Three Ultra-Clear Thinwall centrifuge tubes (5 mL) were used for each OMV sample and the control.

For the membrane fusion assay, OMVs were labeled with the lipid-binding dye octadecyl-rhodamine B chloride (R18) according to Gnopo & Putnam, 2020. OMVs in PBS or PBS alone as a control (2.6 mL) were incubated with 8 µg/ml R18 for 1 h at room temperature. Two variants of the OptiPrep™ density gradient were used: the finer variant with layers from 21% to 28%, and the other variant with layers of 15%, 20%, 25%, then 30% containing R18-OMV or the control (PBS with R18), and the cushion of 35% Optiprep at the bottom. Both gradient variants resulted in complete removal of unbound R18, as the signals were absent in the control, whereas they were observed in the R18-OMV-treated roots at the same time points.

All gradients were centrifuged at +4°C/100,000 g/18h in a SW55Ti rotor, Beckman Coulter - Optima L-80 XP Ultracentrifuge. A ring containing fluorescently labeled OMVs was collected from each of the three tubes and pooled. The same amount of gradient medium was collected from the control at the same position in the tubes. Iodixanol was removed by washing in 10 volumes of PBS by ultrafiltration in an Amicon® Ultra Centrifugal Filter, 100 kDa MWCO, at +4°C/1500g and finally concentrated to 300 µL, containing 10^10^ particles/mL, as determined by NTA (Supplementary files 2, 3 and 4).

### Live Imaging Using Confocal Laser Scanning Microscopy CLSM

Four-day-old tomato or *A. thaliana* seedlings roots were rinsed in PBS and then incubated in DiD- or R18-labeled OMVs or the corresponding controls at room temperature for 1 to 4 hours. After washing three times in PBS, seedlings were observed under CLMS.

Images were captured using a Leica TCS SP8 confocal microscope with 20x or 63x objectives. Z-stacks at 10 µm per slice were acquired using LasAF software (Leica).

## Results

### Isolation and characterization of OMVs secreted by *P. phytofirmans* PsJN

To examine how OMVs produced by plant beneficial rhizospheric bacteria interact with plant cells, we isolated and characterized the OMVs secreted by *P. phytofirmans* PsJN, which is well known for its plant-promoting and protective effects across various host plants. The bacteria were grown in a mineral medium (MSM) to minimize impurities in vesicle preparations. After thoroughly removing bacterial cells and debris, OMVs were isolated and purified using a commercially available affinity-based column. Before further analyses, the buffer was exchanged to PBS through dialysis.

To determine concentration, yield, and size distribution, NTA analysis was performed. As shown in Fig. 1A, particles ranged from 70 to 250 nm, with a median diameter between 110 and 125nm (representative measurement presented in Supplementary file 1). Particle concentration in OMV isolates ranged from 10^9^ to 10^10^ particles/mL. A liter of bacterial culture with an optical density (OD_600_) of 1 yielded up to 10^11^ particles in total. The outer membrane origin of nanoparticles was confirmed by measuring the lipopolysaccharide (LPS) concentration, which was 10^6^ EU per mL of OMV isolate. Protein concentration in the vesicle preparation, measured by the BCA method, was 9 -11 µg/mL, while RNA concentration was 3-4 ng/mL (Fig 1 B).

**Figure 1.**
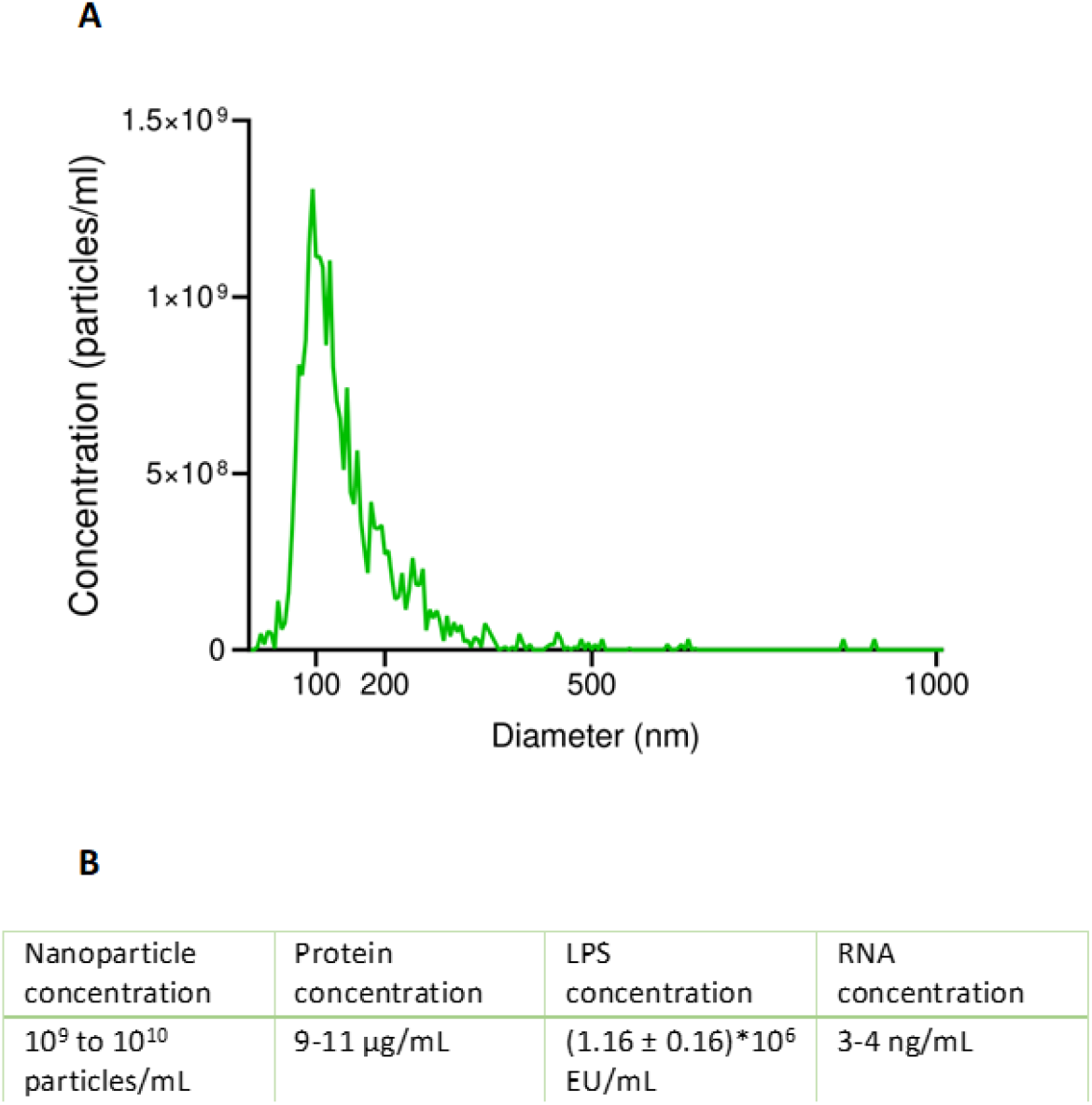
Characterization of OMVs isolated from *P. phytofirmans* PsJN liquid culture. **A)** NTA analysis of particle concentration and size distribution. **B)** Particle, protein, LPS and RNA concentration in PsJN-OMV isolates.

### PsJN-OMVs contact the root epidermal cells

To investigate whether PsJN-OMVs contact root cells of tomato and *A. thaliana*, we labeled OMVs with a lipid-binding dye, Vybrant™ DiD, which efficiently incorporates into biological membranes. The strong affinity of lipophilic dyes for lipids necessitates rigorous controls and thorough washing to prevent unintended labeling of the cell membrane alongside vesicles; therefore, several methods for removing unbound dye were tested (data not shown). Separating DiD-labeled vesicles on a discontinuous density gradient, followed by an ultrafiltration washing step, was chosen as the most reliable method, as the red signals were consistently absent in root cells treated with the corresponding gradient layer of the control (DiD dye in PBS). Contrary, roots of both tomato and *A. thaliana* treated with DiD-labeled OMVs displayed clear red signals. The strongest signals were observed on root hairs (Fig. 2).

**Figure 2.**
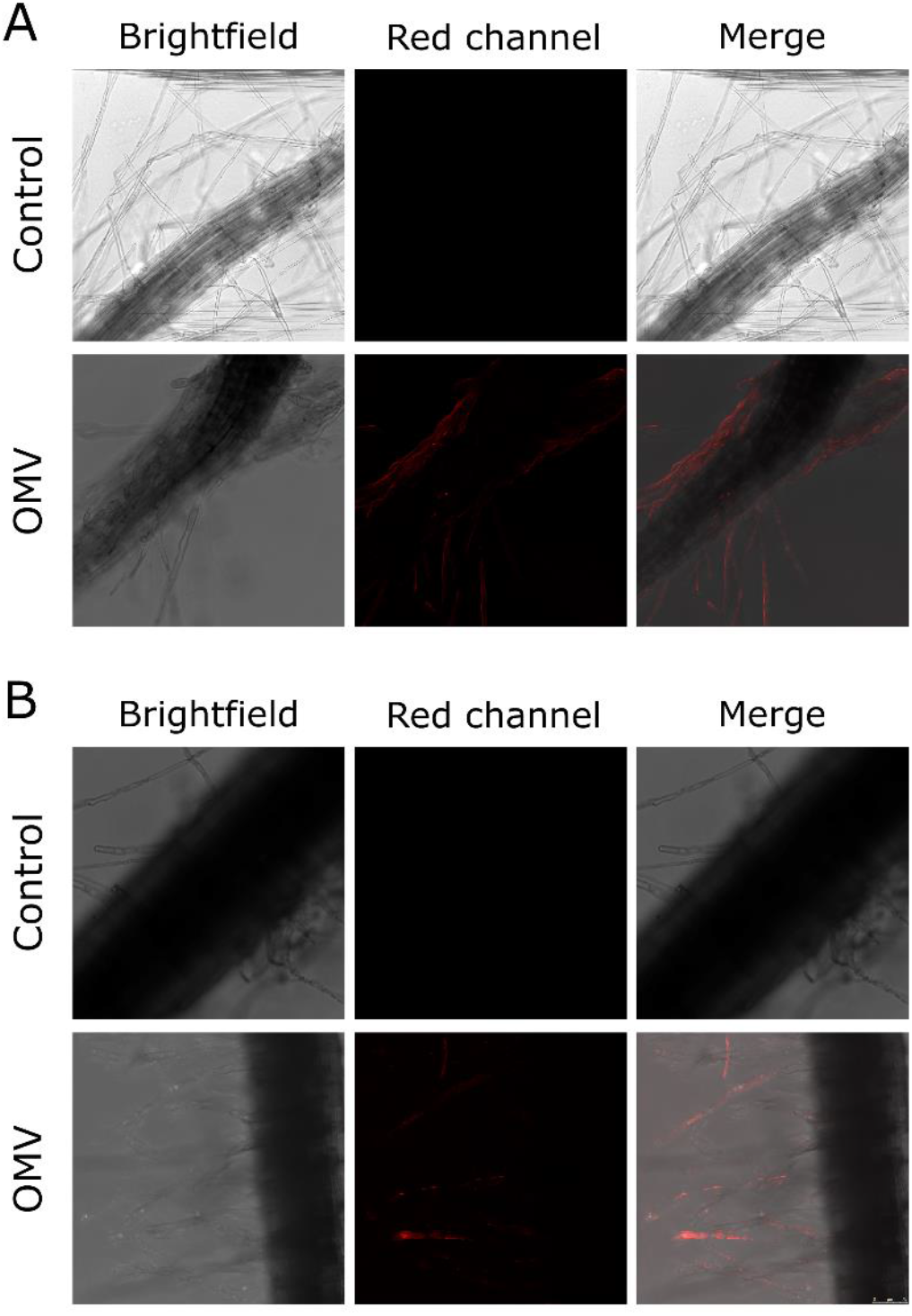
PsJN-OMVs contact root epidermal cells. Roots of four-day old seedlings of **A)** *A. thaliana* or **B)** tomato, were incubated with PsJN-OMVs labelled with Vybrant™ DiD. PBS with the same amount of dye served as a control. The unbound dye was removed from both DiD-labeled OMVs and the control by separation on density gradient and ultrafiltration.

### PsJN-OMVs fuse with the membranes of root epidermal cells

To determine whether PsJN-OMVs fuse with the membranes of root epidermal cells, we used the rhodamine-R18 dye. This lipid-binding dye is self-quenched when present in high concentrations within membranes. However, when the membrane containing R18 fuses with another, unlabeled membrane, the dye diffuses, leading to dequenching and fluorescence detection. After labeling, unbound dye was removed using a density gradient, and R18-labeled OMVs or the appropriate control were applied to the roots of tomato and *A. thaliana* seedlings. Strong red signals were observed in the root epidermal cells of both plants, indicating that OMVs fuse with plant cellular membranes (Fig. 3).

**Figure 3.**
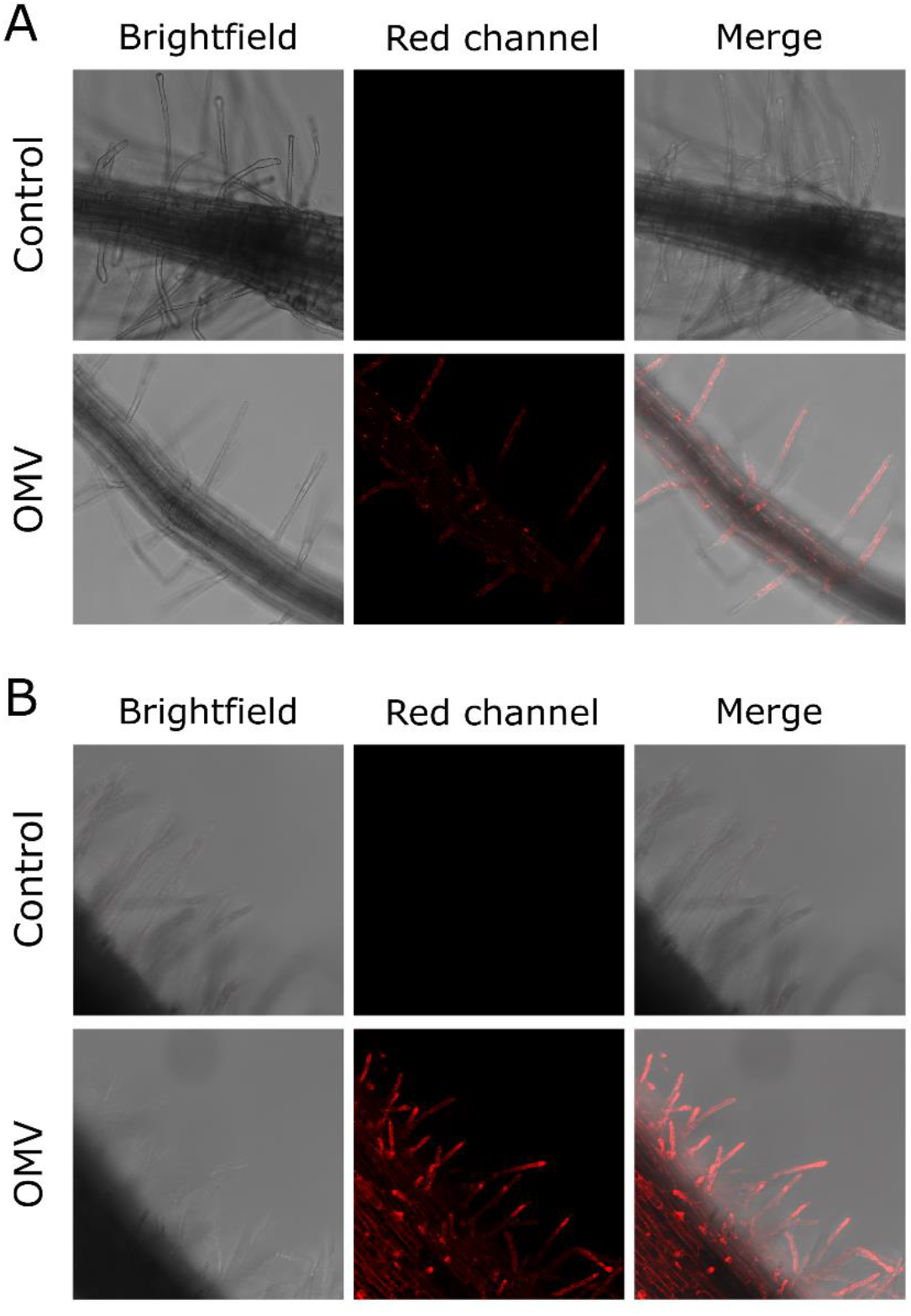
Rhodamine R18-labeled OMV, which fluoresce red upon fusion with the host cell membrane, are found localized in root epidermal cells. Roots of four-day old seedlings of **A)** *A. thaliana* or **B)** tomato, were incubated with PsJN-OMVs labelled with Rhodamine R18. PBS with the same amount of dye served as a control. The unbound dye was removed from both R18-labeled OMVs and the control by separation on density gradient and ultrafiltration.

## Discussion

Bacterial extracellular vesicles are shown to affect eukaryotic cells, thereby participating in bacterial– host interactions and influencing host responses (Kunsmann et al., 2015; Zhao et al., 2025). While OMVs produced by gram-negative human pathogenic and commensal bacteria have received considerable attention lately, the properties and role of OMVs from plant-associated bacteria have been very poorly investigated. For a long time, the exchange of EVs between plant cells and their environment was in doubt, since the pores in the plant cell wall are much smaller than the diameter of EVs (Carpita et al., 1979). However, cell wall-degrading enzymes, such as cellulase, xylanase, pectin lyase, and lipases, were found associated with bacterial vesicles (Feitosa-Junior et al., 2019; Sidhu et al., 2008). The release of these components may facilitate the loosening of the cell wall, allowing OMVs to pass through the cell wall matrix (Tran et al., 2022). OMVs produced by both pathogenic and beneficial bacteria carry immunogenic molecules and can trigger immune responses in plants, priming them against bacterial and fungal infections (Bahar et al., 2016; Janda et al., 2023; McMillan et al., 2021). On the other hand, OMVs produced by some phytopathogens exert immunosuppressive effects, facilitating pathogen invasion (Wu et al., 2024). Other ways in which OMVs’ action supports the spread of pathogens were also observed, e.g., by blocking bacterial attachment to surfaces, thereby increasing bacterial movement within the xylem, and facilitating colonization of the plant (Ionescu et al., 2014). Although these effects highlight the important role of OMVs in bacterial-plant interactions, very few studies have explored how OMVs convey signals and deliver effector molecules. The only thorough investigation of this subject was done by Tran et al. (2022), who demonstrated integration of pathogenic bacterium *Xanthomonas campestris* OMVs into the plasma membrane of *A. thaliana* cells.

Regarding beneficial bacteria, the mechanism by which OMVs interact with plant cells remains elusive. Here, we demonstrate that OMVs produced by the rhizospheric bacterium *P. phytofirmans* PsJN directly interact with epidermal root cells. First, this was observed after incubating *A. thaliana* and tomato roots with OMVs labeled with the lipid-binding dye Vybrant^TM^ DiD. To determine whether OMVs were present only on the cell surface or if they contacted and fused with the membranes of epidermal cells, we employed the rhodamine-R18 dye, which exhibits fluorescence only upon membrane fusion (Bomberger et al., 2009). Strong red signals observed in root epidermal cells indicate that OMVs’ membranes indeed fused with plant cell membranes.

Some previous studies have suggested that OMVs of beneficial bacteria directly contact plant cells and deliver their contents onto the plant cell surface or into the plant cells during the establishment or maintenance of symbiosis. Nod factors, bacterial signal molecules essential for the onset of rhizobia-legume symbiosis, are found in isolates of OMVs produced by *Rhizobium etli* CE3 grown in the presence of flavonoid naringenin, i.e., in symbiosis-mimicking conditions (Taboada et al., 2019). The authors suggested that OMVs can serve as a transport vehicle for Nod factors, protecting them from degradation in the environment and enabling their application to the root epidermal cells in a concentrated form. OMVs produced by rhizobia in symbiosis-mimicking conditions caused notable deformation of root hairs, increased expression of nodulation genes, and decreased expression of defense genes (D. Li et al., 2022; Taboada et al., 2019). To date, direct contact of rhizobial OMVs with the plant cell surface has not been observed, and the mechanism of the interaction of bacterial vesicles with legume root hairs has not been resolved.

Our results indicate that OMVs produced by rhizospheric non-rhizobial plant beneficial bacteria directly contact root cells. Additionally, we observed the fusion of OMVs’ membranes with plant cell membranes, which can include the plasma membrane or membranes of some intracellular compartments after the endocytosis of the vesicles. The fusion of pathogenic OMVs with the plasma membrane of the eukaryotic host cells has been previously observed in human and plant cells (Rompikuntal et al., 2012; Tran et al., 2022). If R18 signals that we observed originate from PsJN-OMVs fusion with the plasma membranes of plant cells, OMVs would deliver their content into the epidermal cell cytoplasm, while their membranes would become integrated into the plasma membrane. The integration event itself is shown to modify membrane lipid packing, altering its physicochemical properties and consequently impacting cellular processes (Tran et al., 2022). The authors suggested that this mechanism plays a crucial role in microbe–plant cell interactions independently of MAMPs (microbe-associated molecular patterns). The saturation level of lipids is an important factor influencing membrane rigidity and fluidity. Plant plasma membranes are enriched in unsaturated fatty acids to maintain the fluidity required for plant cell functions (Reszczyńska & Hanaka, 2020). On the other hand, the lipid saturation level in OMVs is substantially higher, contributing to increased membrane rigidity, which is necessary for vesicle formation and stability (Tashiro et al., 2011). Therefore, the insertion of OMVs alters the saturation and symmetry levels of phospholipids in the plant plasma membrane, resulting in increased lipid order and packing density (Tran et al., 2022). The enhancement of lipid order and compartmentalization of membranes is considered to play a crucial role in refining signaling scaffolds on the plant plasma membrane in response to both abiotic and biotic environmental stimuli (Huang et al., 2019; Ke et al., 2021; Ma et al., 2021; Tran et al., 2020). Plant surface nanodomains with a high degree of order are crucial for immune signaling during interactions between plants and pathogens (Bücherl et al., 2017; Huang et al., 2019). Tran et al. (2022) demonstrated that the integration of OMVs in the plant plasma membrane is dependent on phytosterol- and remorin-containing nanodomains in the recipient membrane. On the other hand, the integration of OMVs induces the clustering of remorin proteins and enhances nanodomain assembly, establishing a feedback loop that leads to increased compartmentalization of the plasma membrane and, eventually, priming of plant immunity. Additionally, the integration of OMVs into the plasma membrane increased the endocytosis rate; nonetheless, OMVs were not transported into the cells via endocytosis. Instead, their membranes remained inserted in the plasma membrane, with restricted exchange with the surrounding lipids of the recipient membrane.

However, aside from vesicle fusion with the plasma membrane, there are other possible routes for OMVs’ entrance and cargo delivery to the recipient cell, including clathrin-dependent or independent endocytosis (O’Donoghue & Krachler, 2016). The uptake route depends on OMVs’ composition and surface properties, which differ between species or strains and change under the influence of environmental cues and signals from interacting organisms (McBroom & Kuehn, 2007; O’Donoghue & Krachler, 2016; Taboada et al., 2019). Additionally, vesicles produced by the same cells under the same conditions are not homogeneous, but vary in size and characteristics (Ñahui Palomino et al., 2021). If PsJN-OMVs are internalized into plant cells by endocytosis, the observed R18 signals would then originate from the subsequent OMVs membrane fusion with the membranes of some intracellular compartment. This way of OMVs’ entry would also lead to the delivery of the OMVs’ cargo and its interaction with plant cellular components. For instance, Cañas et al. (2016) demonstrated that R18-labeled OMVs, originating from probiotic and commensal *E. coli* strains, entered human intestinal epithelial cells *via* clathrin-mediated endocytosis. This entry route was also shown for OMVs of pathogenic bacteria (Parker et al., 2010). Another possibility for the endocytic internalization of OMVs is *via* clathrin-independent endocytosis, as demonstrated for the interaction between periodontal pathogen OMVs and human cells (Furuta et al., 2009).

At this point, it is difficult to assume whether cargo carried by non-rhizobial plant beneficial OMVs could trigger signaling events in root cells, and if so, which molecules would be involved. The recognition of non-rhizobial PGPB likely involves overlapping symbiosis and immunity pathways, with plants balancing positive and antagonistic signals to determine compatibility and mutual benefit. Unlike rhizobia-legume symbiosis, where Nod factors and LysM receptors are well-defined, signaling mechanisms underlying mutualistic interactions between plants and non-rhizobial beneficial bacteria are still being unraveled.

### Conclusion and future perspectives

Herein, we have demonstrated that OMVs produced by PGPB *P. phytofirmans* PsJN directly contact roots of tomato and *A. thaliana* plants and fuse with the membranes of their epidermal cells. Membrane fusion was observed using the rhodamine R18 dye, and this event can occur at the plasma membrane or within the membranes of intracellular compartments. Both scenarios would lead to alterations in the recipient membrane properties and the delivery of OMVs’ cargo to plant cells. To the best of our knowledge, our study is the first to report direct interaction of OMVs produced by plant-beneficial bacteria with plant cells.

The colonization of plant roots by beneficial rhizospheric bacteria involves a multi-stage process driven by dynamic chemical communication between the plant and microbes. Interactions with beneficial or pathogenic microbes trigger distinct immune and metabolic responses in plant roots, ultimately leading to either the promotion or suppression of microbial colonization. Whether and how OMVs participate in this complex interplay is an open and intriguing question. The findings presented in our study lay the foundation for future research on the potential role of OMVs in the early stages of root colonization and the establishment of symbiotic relationships. While the biomedical potential of OMVs has been recognized over the past two decades, and extensive efforts have been made to utilize microbial vesicles for therapeutic purposes, many aspects of OMVs’ role in plant–microbe interactions remain unclear. Therefore, increasing our understanding in this area could enable the use of these nano-vehicles to develop eco-friendly solutions for overcoming environmental challenges and supporting sustainable agriculture.

## Supplementary material

<u>Supplementary file 1</u>. Representative NTA analysis of nanoparticles in PsJN-OMV isolate

<u>Supplementary file 2</u>. Graph representing the size distribution of nanoparticles in R18-labeled PsJN-OMV sample or the appropriate control (PBS with the same concentration of added R18 dye), after separation on a density gradient and ultrafiltration purification

<u>Supplementary file 3</u>. Representative NTA analysis of nanoparticles in R18-labeled PsJN-OMV sample, after separation on density gradient and ultrafiltration purification

<u>Supplementary file 4</u>. Representative NTA analysis of nanoparticles in the control (PBS with the same concentration of added R18 dye as in the OMV sample), after separation on a density gradient and ultrafiltration purification

## Acknowledgements

We are very grateful to our colleagues Marija Schwirtlich, Mila Ljujić, and Jovana Jasnić, UB IMGGE, for selfless help and support in CLSM microscopy. We are also thankful to Aleksandra Korać, Center for Electron Microscopy, UB Faculty of Biology, and Marija Milivojević, UB Department of Inorganic Chemical Technology.

