## Supplementary File 1 for "Outer Membrane Vesicles (OMVs) Produced by Plant Beneficial Rhizospheric Bacteria Enter Root Epidermal Cells"

PARTICLE METRIX

### ZETAVIEW®

www.particle-metrix.com

Electrophoresis & Brownian Motion
Video Analysis
Laser Scattering Microscopy

Video Operator: ZetaUser

Operator (Report): ZetaUser

**Sample Parameters**
Sample Name: OMV-12-6-25
Sample Info 1:
Sample Info 2:
Sample Info 3:
Electrolyte:
Temperature: 25.05 °C sensed
pH 7.0 entered
Conductivity: 15000.00 µS/cm sensed

**Instrument Parameters**
Laser Wavelength: 488 nm
Filter Wavelength: Scatter
Sensitivity: 78
Shutter: 100

**SOP: MK_ssc**
Size Distribution 1 Cycle 11 Positions
Description:

**Result (sizes in nm)**

| | Number | Concentration | Volume |
|---|---|---|---|
| Median (X50) | 113.7 | 113.7 | 297.7 |
| StdDev | 79.3 | 79.2 | 182.5 |

Concentration: 4.0E+7 Particles / mL
Dilution Factor: 250
Original Concentration: 9.9E+9 Particles / mL

**Quality**
Average Counted Particles per Frame: 91
Number of Traced Particles: 427
1 Positions Removed for Analysis

**Analysis Parameters**
Max Area 1000, Min Area 10, Min Brightness 25, Classes/Decade: 64

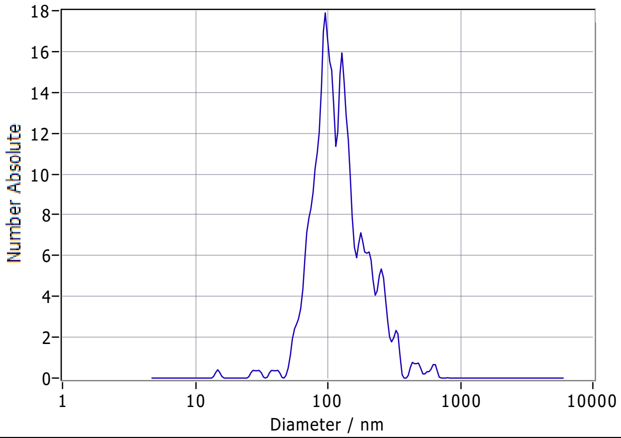


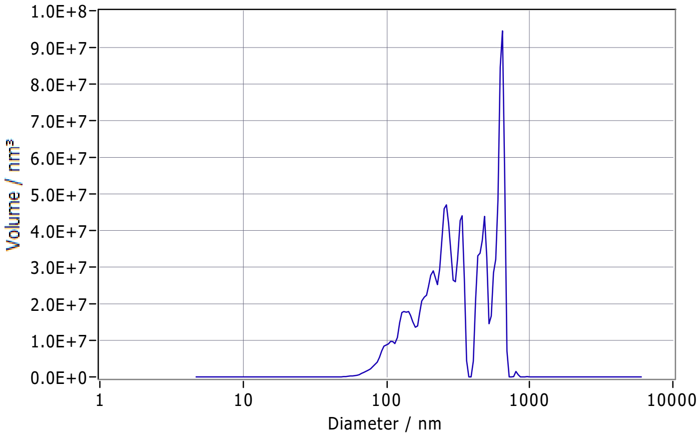


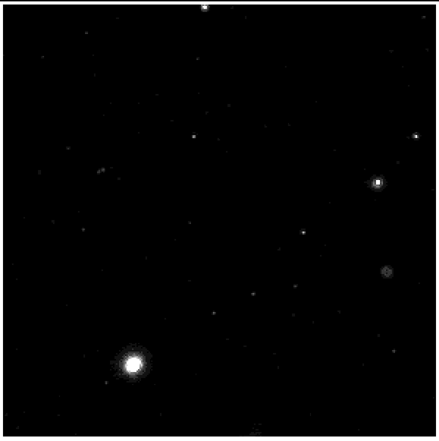

**Peak Analysis (Concentration)**

| Diameter / nm | Number Absolute | FWHM / nm | Percentage |
|---|---|---|---|
| 96.7 | 1.8E+1 | 34.1 | 48.1 |
| 128.6 | 1.6E+1 | 29.9 | 28.9 |
| 178.2 | 7.1E+0 | 42.8 | 12.5 |
| 252.9 | 5.3E+0 | 38.7 | 6.6 |
| 324.1 | 2.3E+0 | 47.2 | 3.9 |

**X Values** (all sizes are given in nm)

| | Number | Concentration | Volume |
|---|---|---|---|
| X10 | 71.1 | 71.1 | 132.4 |
| X50 | 113.7 | 113.7 | 297.7 |
| X90 | 232.9 | 232.9 | 623.9 |
| Span | 1.4 | 1.4 | 1.7 |
| Mean | 138.2 | 138.2 | 354.8 |
| StdDev | 79.3 | 79.2 | 182.5 |

**Comment**

(Signature)

Analyzed Video: C:\Users\ZetaView\Documents\ZetaView Data\Rezultati\2025-06-16 Majad\20250616_0004_OMV-12-6-25_size_488.avi

Experiment: 2025-06-16 09:24 ZetaView S/N 22-846, Software ZetaView (version 8.05.16 SP3)

Report: 2025-06-16 09:24 Software ZetaView (version 8.05.16 SP3)
