## Supplementary File 3 for "Outer Membrane Vesicles (OMVs) Produced by Plant Beneficial Rhizospheric Bacteria Enter Root Epidermal Cells"

### ZETAVIEW®

www.particle-metrix.com

Electrophoresis & Brownian Motion
Video Analysis
Laser Scattering Microscopy

Video Operator: ZetaUser

Operator (Report): ZetaUser

**Sample Parameters**
Sample Name: OMV-13-6-25
Sample Info 1:
Sample Info 2:
Sample Info 3:
Electrolyte:
Temperature: 25.00 °C sensed
pH 7.0 entered
Conductivity: 15000.00 µS/cm sensed

**Instrument Parameters**
Laser Wavelength: 488 nm
Filter Wavelength: Scatter
Sensitivity: 78
Shutter: 100

**SOP: MK_ssc**
Size Distribution 1 Cycle 11 Positions
Description:

**Result (sizes in nm)**

| | Number | Concentration | Volume |
|---|---|---|---|
| Median (X50) | 129.1 | 129.1 | 349.6 |
| StdDev | 88.5 | 88.3 | 263.4 |

Concentration: 2.0E+7 Particles / mL
Dilution Factor: 500
Original Concentration: 9.9E+9 Particles / mL

**Quality**
Average Counted Particles per Frame: 45
Number of Traced Particles: 260

**Analysis Parameters**
Max Area 1000, Min Area 10, Min Brightness 25, Classes/Decade: 64

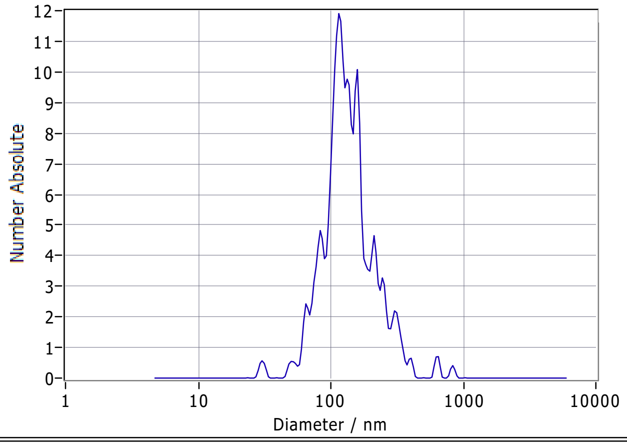


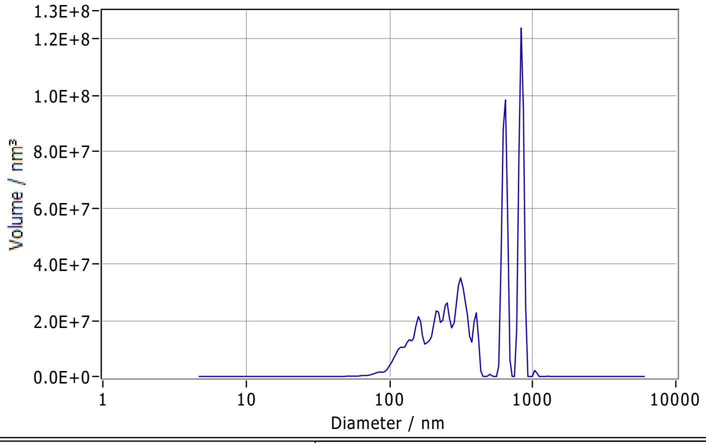


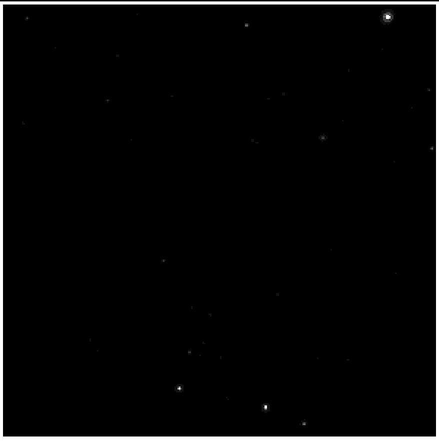

**Peak Analysis (Concentration)**

| Diameter / nm | Number Absolute | FWHM / nm | Percentage |
|---|---|---|---|
| 115.7 | 1.2E+1 | 35.7 | 44.8 |
| 157.0 | 9.8E+0 | 25.1 | 21.9 |
| 211.3 | 4.6E+0 | 37.2 | 14.4 |
| 83.8 | 4.8E+0 | 25.5 | 12.7 |
| 306.1 | 2.2E+0 | 53.3 | 6.2 |

**X Values** (all sizes are given in nm)

| | Number | Concentration | Volume |
|---|---|---|---|
| X10 | 80.3 | 80.3 | 149.1 |
| X50 | 129.1 | 129.1 | 349.6 |
| X90 | 241.6 | 241.6 | 820.3 |
| Span | 1.2 | 1.2 | 1.9 |
| Mean | 152.2 | 152.2 | 464.5 |
| StdDev | 88.5 | 88.3 | 263.4 |

**Comment**

(Signature)

Analyzed Video: C:\Users\ZetaView\Documents\ZetaView Data\Rezultati\2025-06-16 Majad\20250616_0006_OMV-13-6-25_size_488.avi

Experiment: 2025-06-16 09:31 ZetaView S/N 22-846, Software ZetaView (version 8.05.16 SP3)

Report: 2025-06-16 09:32 Software ZetaView (version 8.05.16 SP3)
