## Supplementary File 4 for "Outer Membrane Vesicles (OMVs) Produced by Plant Beneficial Rhizospheric Bacteria Enter Root Epidermal Cells"

**ZETAVIEW®**

www.particle-metrix.com

Electrophoresis & Brownian Motion
Video Analysis
Laser Scattering Microscopy

Video Operator: ZetaUser

Operator (Report): ZetaUser

**Sample Parameters**
Sample Name: K-za-13-6-25
Sample Info 1:
Sample Info 2:
Sample Info 3:
Electrolyte:
Temperature: 24.98 °C sensed
pH 7.0 entered
Conductivity: 13889.00 µS/cm sensed

**Instrument Parameters**
Laser Wavelength: 488 nm
Filter Wavelength: Scatter
Sensitivity: 78
Shutter: 100

**SOP: MK_ssc**
Size Distribution 1 Cycle 11 Positions
Description:

**Result (sizes in nm)**

| | Number | Concentration | Volume |
|---|---|---|---|
| Median (X50) | 138.3 | 138.3 | 268.7 |
| StdDev | 77.6 | 77.5 | 138.8 |

Concentration: 4.4E+7 Particles / mL
Dilution Factor: 20
Original Concentration: 8.9E+8 Particles / mL

**Quality**
Average Counted Particles per Frame: 101
Number of Traced Particles: 587
1 Positions Removed for Analysis

**Analysis Parameters**
Max Area 1000, Min Area 10, Min Brightness 25, Classes/Decade: 64

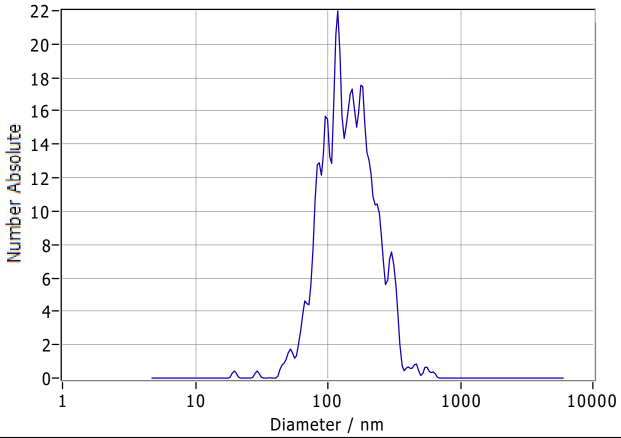


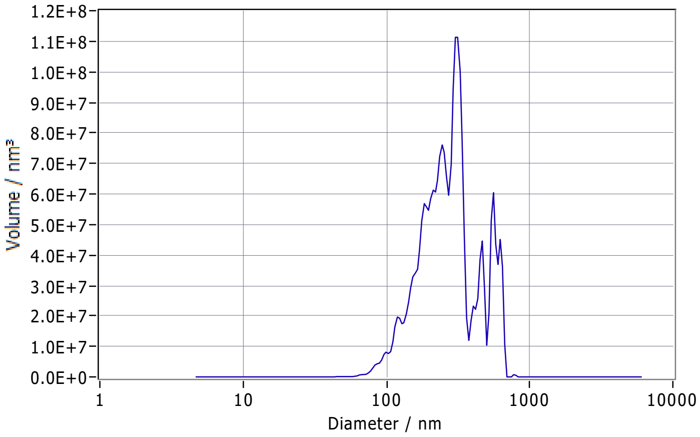


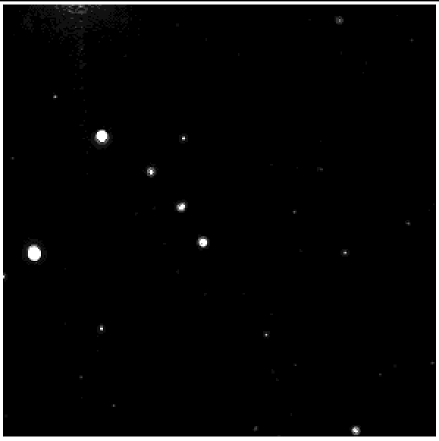

**Peak Analysis (Concentration)**

| Diameter / nm | Number Absolute | FWHM / nm | Percentage |
|---|---|---|---|
| 178.6 | 1.8E+1 | 56.9 | 33.8 |
| 119.0 | 2.2E+1 | 13.9 | 25.5 |
| 98.5 | 1.6E+1 | 40.3 | 21.4 |
| 151.4 | 1.7E+1 | 15.7 | 11.2 |
| 302.8 | 7.5E+0 | 56.0 | 8.1 |

**X Values** (all sizes are given in nm)

| | Number | Concentration | Volume |
|---|---|---|---|
| X10 | 80.8 | 80.8 | 148.4 |
| X50 | 138.3 | 138.3 | 268.7 |
| X90 | 255.6 | 255.6 | 537.6 |
| Span | 1.3 | 1.3 | 1.4 |
| Mean | 158.7 | 158.7 | 301.1 |
| StdDev | 77.6 | 77.5 | 138.8 |

**Comment**

(Signature)

Analyzed Video: C:\Users\ZetaView\Documents\ZetaView Data\Rezultati\2025-06-16 Majad\20250616_0013_K-za-13-6-25_size_488.avi

Experiment: 2025-06-16 10:34 ZetaView S/N 22-846, Software ZetaView (version 8.05.16 SP3)

Report: 2025-06-16 10:35 Software ZetaView (version 8.05.16 SP3)
