## Supplementary figures and images for "Outer Membrane Vesicles (OMVs) Produced by Plant Beneficial Rhizospheric Bacteria Enter Root Epidermal Cells"

### Supplementary File 2

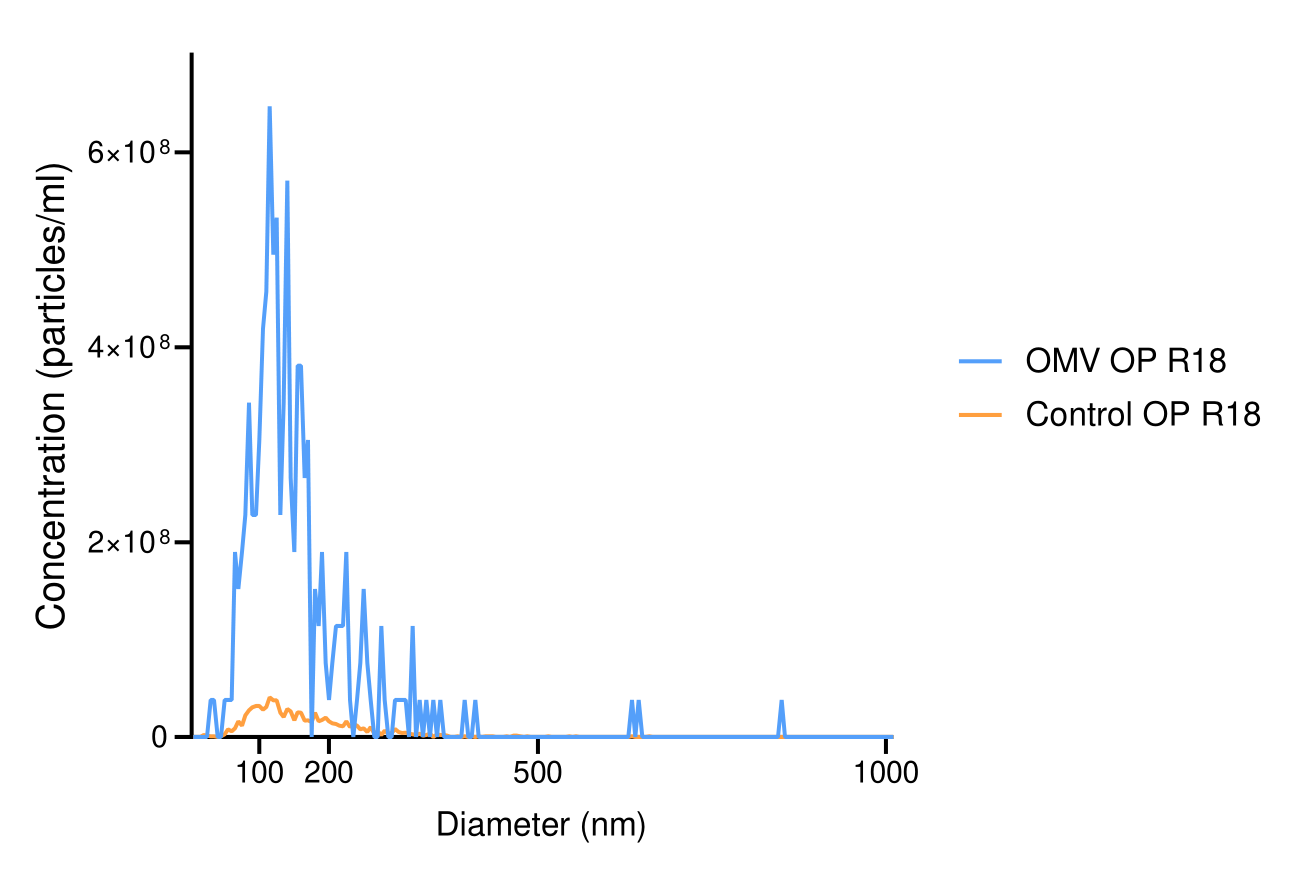
